# Molecular insights on the mechanism of *α*_1_-antitrypsin condensate formation and maturation

**DOI:** 10.1101/2025.03.11.642656

**Authors:** Ignacio Sanchez-Burgos, Andres R. Tejedor, Rosana Collepardo-Guevara, Jorge Bernardino de la Serna, Jorge R. Espinosa

## Abstract

The deficiency of *α*_1_-antitrypsin protein is a genetic disorder characterized by the accumulation of misfolded protein aggregates within hepatocytes, leading to liver dysfunction. In the lung, it is found in macrophages, bronchial and epithelial alveolar cells type 2, leading to pulmonary emphysema. Despite extensive research, the precise mechanism underlying the formation of *α*_1_-antitrypsin inclusion bodies remain elusive. In this study, we combine equilibrium and non-equilibrium molecular dynamics simulations to elucidate the intricate process of *α*_1_-antitrypsin condensate formation and maturation. Our mechanistic model explains cluster accumulation—specifically the onset of this pathogenesis—through the emergence of phase-separated liquid-like protein droplets, which subsequently undergo inter-protein *β*-sheet transitions between misfolded variants, resulting in solid-like clusters. We find that this mechanism only applies to the misfolded variant, Z-*α*_1_-antitrypsin, which phase-separates driven by its disordered C-terminus. In contrast, the native protein, M-*α*_1_-antitrypsin, shows much lower propensity to phase-separate and later form kinetically trapped aggregates. Furthermore, we explore how Z-*α*_1_-antitrypsin exhibits an increased capacity to form condensates near external walls with different types of interactions. Such conditions can be similar to those found within the endoplasmic reticulum membrane, where phase separation and hardening take place. Overall, our results shed light on the molecular basis of *α*_1_-antitrypsin-related disorders and provide valuable microscopic insights for the development of therapeutic strategies targeting protein misfolding and aggregation-related disorders.

**Author summary:** Alpha-1 antitrypsin deficiency is a genetic disorder that can cause liver and lung disease. It occurs when the protein *α*_1_-antitrypsin folds incorrectly and accumulates in harmful clumps inside cells. Exactly how these clumps form has been unclear. In this study, we used computer simulations to reveal the steps behind this process. We found that the faulty version of the protein first collects into liquid-like droplets that later harden into solid clusters. This behavior is driven by a flexible tail region of the misfolded protein. In contrast, the normal version is far less likely to clump. Our findings provide new insight into disease onset and may guide the development of therapies.

## I. INTRODUCTION

In living organisms, there is a crucial regulatory mechanism responsible for maintaining the correct balance of cellular proteins, known as proteostasis^1^. Disruptions in this system have been linked to the development of several common human diseases^2^ such as cystic fibrosis^3^ or type 2 diabetes^4^. At the heart of this regulatory network lies the intricate process of protein folding, where newly synthesized polypeptide sequences must fold correctly to attain their functional three-dimensional structure^5,6^. It is widely recognized that protein folding can occur during the synthesis process, which aids in the efficient achievement of functional structure. However, protein folding is also susceptible to errors, as evidenced by studies showing that misfolding can occur during protein synthesis, leading to degradation of aberrantly folded sequences^7–9^.

*α*_1_-antitrypsin, a glycoprotein crucial for regulating protease activity in the human body^10^, is implicated in the pathology of *α*_1_-antitrypsin deficiency (AATD), a genetic disorder^11^ that leads to liver dysfunction and pulmonary emphysema. This protein is synthesized within hepatocytes and undergoes folding in the endoplasmic reticulum before being secreted into the bloodstream, where it primarily functions as an inhibitor of neutrophil elastase in the lungs^10,11^. While its correct folding has been shown to be essential for its functional activity^12^, mutations in the *α*_1_-antitrypsin gene lead to incorrect folding of the sequence, resulting in the formation of insoluble bodies within hepatocytes, which leads to AATD pathogenesis^11,13–15^. To elucidate the mechanisms underlying *α*_1_-antitrypsin misfolding, researchers have conducted detailed investigations into the folding dynamics of nascent polypeptide sequences as they emerge from the ribosome during the synthesis. These studies have revealed that nascent *α*_1_-antitrypsin can exhibit different folding behaviors in the ribosome, forming intermediates with molten-globule characteristics, where the C-terminus appears to be misfolded^15–17^. While structural analyses have provided major insights into the differences in folding kinetics between wild-type and mutant *α*_1_-antitrypsin variants^11,18^, shedding light on the precise origin of misfolding and their implications in health and disease remains a challenge.

The wild-type *α*_1_-antitrypsin sequence, termed M-*α*_1_-antitrypsin, rapidly folds and carries out the biological function of regulating the activity of proteases^12,15^. However, the mutated Z-*α*_1_-antitrypsin variant contains a glutamic acid to lysine mutation in the C-terminal region (E342K), which impairs the protein’s ability to attain its functional structure^17^. This variant has been identified as the principal cause of *α*_1_-antitrypsin deficiency (AATD)^11,13,14,17^. In the original protein sequence, Glu_342_ forms a salt bridge to Lys_290_, which is not present in the Z variant, and is thought to be the reason behind the delay in the sequence folding^14^. Since aberrant aggregates formed by the Z variant of *α*_1_-antitrypsin resemble those observed in protein condensate-related neurodegenerative disorders^19–22^, here we apply a computational approach developed by us to explore the protein phase behavior at submolecular level from individual sequences to higher-order assemblies^23–25^.

We perform Molecular Dynamics (MD) simulations with a sequence-dependent coarse-grained protein model^26^ to evaluate the phase diagrams of the M and Z variants, providing insights into their thermodynamic coexistence lines and intermolecular liquid network connectivity. Unlike other commonly used computational approaches for simulating protein structural dynamics and phase behaviour—such as all-atom force fields (CHARMM^27^, AMBER^28^) or high-resolution coarse-grained models like MARTINI^29^—our model does not rely on explicit solvent definition. While these well-established models can capture atomic detail, their high computational cost limits system size and simulation timescales, making them poorly suited for mesoscale condensate dynamics. In contrast, the chosen sequence-dependent model, CALVADOS2^26^, uses an implicit solvent and reduced resolution of 1-bead per amino acid, allowing simulations of hundreds of protein replicas and capturing condensate formation, and disorder-to-order transitions within feasible simulation timescales^30^. Furthermore, to decipher the clustering mechanism of Z-*α*_1_-antitrypsin, we model how different protein replicas can dynamically establish inter-protein *β*-sheet transitions within condensates, which over time drive a shift from liquid-like to more solid-like insoluble states similar to those observed in AATD. We characterize the viscosity of these protein assemblies, before and after the formation of intermolecular cross-*β*-sheet structures, and we investigate the interplay between intra- and inter-protein structural changes, which is crucial for understanding the competition between correct folding and the formation of potentially insoluble bodies. Lastly, we simulate how the presence of a surface regulates the propensity of *α*_1_-antitrypsin to form condensates via liquid-liquid phase separation (LLPS). Overall, our comprehensive computational approach provides valuable information on the phase behavior, viscosity, and maturation mechanisms of the M and Z-*α*_1_-antitrypsin protein variants at physiological conditions.

## II. METHODS

To model the different *α*_1_-antitrypsin variants, we employ the residue-level coarse-grained force field CALVADOS2^26^. Within this implicit-solvent model, proteins are simulated as polymers in which each bead represents a given amino acid with its own chemical identity. The CALVADOS2 model accurately predicts the conformational properties of intrinsically disordered proteins and propensities to undergo LLPS for diverse sequences and solution conditions. The intermolecular potential describing the interactions between all pairs of amino acids in the CALVADOS2 is detailed in the Supplementary Material (SM) Section I. To account for the structured domains of the protein, we preserve the relative positions of the residues (centered on the C_*α*_s of each amino acid) that belong to the experimentally reported structured domains (PDB code: 3NE4 for both M and Z variants) with a rigid body integrator. In contrast, intrinsically disordered regions are simulated as flexible polymers using a harmonic bond potential. We use the LAMMPS MD package^31^ to perform our simulations. We use the mature form (with the N-terminus truncated), which is the functional protein that is secreted and active in the plasma^**?**^ . Moreover, to mimic the emergence of inter-protein structured domains, we employ our previously developed aging algorithm^23,24,30^, which dynamically evaluates high-density local fluctuations of protein domains (i.e., low-complexity aromatic-rich segments; LARKS^20,32^) which are prone to develop inter-protein *β*-sheet structures. Under these conditions, disorder-to-order structural transitions are susceptible to occur, and the intermolecular interactions of the involved residues are scaled, accounting for the stronger interaction associated with the formation of inter-protein *β*-sheets^25,33,34^ (further numerical details on the algorithm and the precise simulation technical details and system sizes are described in the SM Section SII).

## III. RESULTS

### A. Phase separation propensity of M and Z-*α*_1_-antitrypsin variants

Recent studies have suggested that the C-terminal region of the Z-*α*_1_-antitrypsin variant exhibits increased conformational flexibility and reduced structural stability, behaving more like a partially disordered or transiently unfolded segment^17,35,36^, in contrast to the native M-*α*_1_-antitrypsin, in which the C-terminus is stably folded into multiple intramolecular *β*-sheet structures. We have examined the structures predicted by AlphaFold^37^ for both the M and Z variants of *α*_1_-antitrypsin, and as expected, AlphaFold predicts virtually identical folded structures for the two variants, consistent with its focus on the most probable equilibrium conformation. This is in agreement with previous structural studies showing that the Z mutation might not alter the final folded state, but instead increase the kinetic barrier to folding and promote transiently unstructured intermediates^36^. Moreover, AlphaFold can be limited in structural prediction of single point mutations as recently disclaimed in the software^37^.

Our working hypothesis is based on the fact that the enhanced conformational flexibility of the Z variant may facilitate the formation of dense, dynamic assemblies as those that drive liquid–liquid phase separation of proteins and form condensates. These condensates could then act as precursors to more ordered solid-like structures. While Ref.^17^ primarily addresses the liquid-to-solid transition, it supports the broader concept that condensate formation can precede structural reorganization into solid-like assemblies. Therefore, our coarse-grained simulations aim to reveal the comparative tendency of the Z and M variants to form such dense, dynamic clusters in the first instance. In Figure 1(a), we sketch the different structural motifs of *α*_1_-antitrypsin: (i) intrinsically disordered regions depicted by straight lines; (ii) *α*-helix structures by ovals; and (iii) *β*-sheet domains indicated by arrows. Dashed lines show the regions of the sequence that rapidly fold in the M variant, while remaining disordered in the Z variant. As can be seen, as a consequence of the (E342K) mutation, there is a segment of the sequence that, if not properly folded, may have the potential to promote phase-separation, and as a consequence, form inter-protein *β*-sheets instead of intramolecular folding. This domain (highlighted with dashed lines in Figure 1(a), and visibly disordered as indicated in Figure 1(b)) could be susceptible to promote LLPS among misfolded variants, and may explain the physicochemical impact of such mutation in the protein phase behavior leading to AATD.

**Figure 1:**
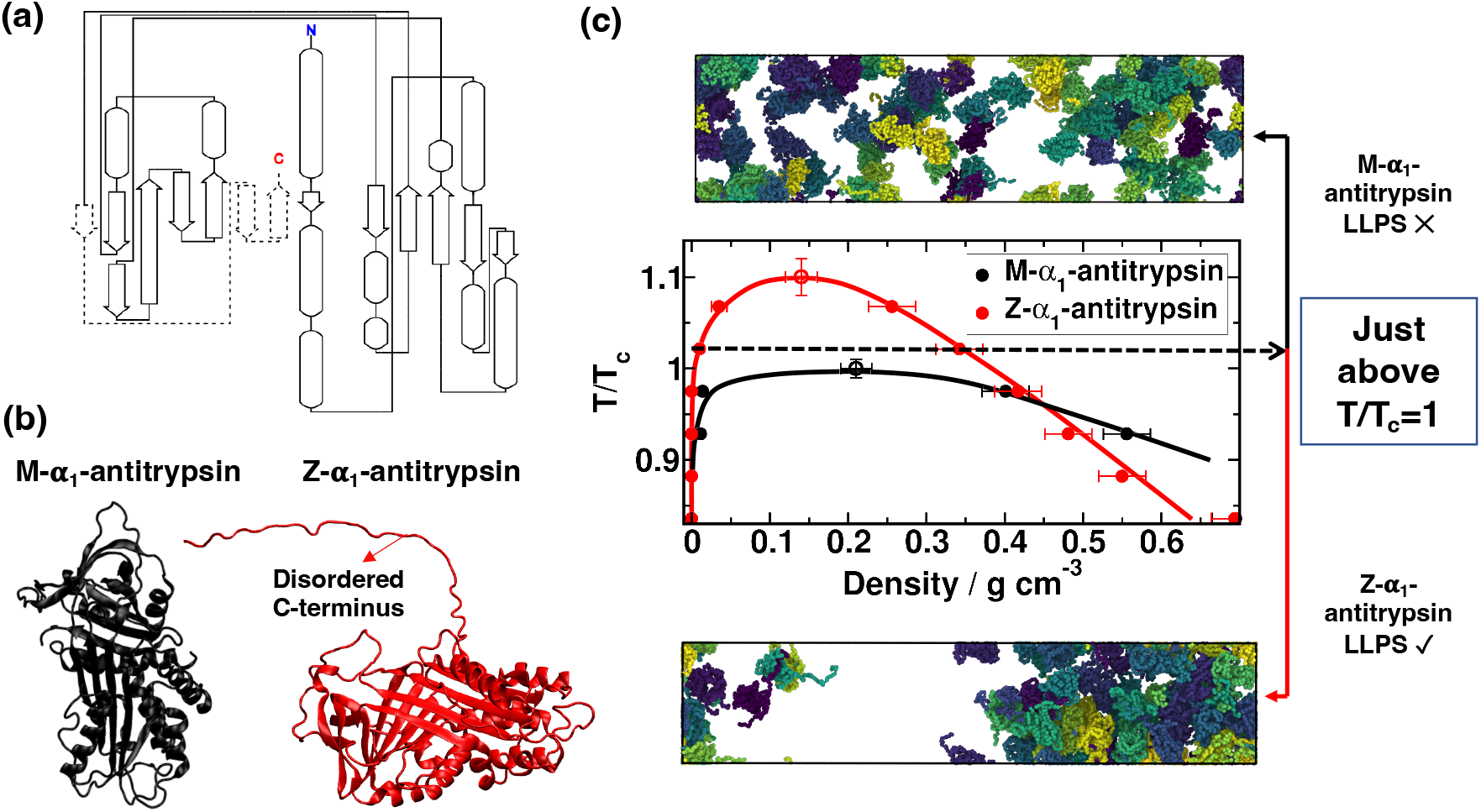
(a) Full sequence representation of *α*_1_-antitrypsin illustrating the intrinsically disordered regions with straight lines, the *α*-helix structures with ovals, and *β*-sheets with arrows. The dashed region has no secondary structure in Z-*α*_1_-antitrypsin. (b) Rendered images of the M and Z variants of *α*_1_-antitrypsin employed in our simulations. (c) Phase diagram in the temperature (normalized by the critical temperature of the M variant) density plane for the M and Z *α*_1_-antitrypsin sequences, along with representative snapshots at T/T_*c*_ ∼1.03, where Z-*α*_1_-antitrypsin displays a condensate coexisting with a diluted phase (Bottom), while M-*α*_1_-antitrypsin cannot undergo phase-separation (Top). The errors are obtained from block analysis to the condensed phase density.

To test this hypothesis, we build both variants using the high-resolution crystal structure (1.8 Å; PDB code^38^: 3NE4) resolved for M-*α*_1_-antitrypsin. We employ the CALVADOS2^26^ force field to model the proteins, and the LAMMPS rigid body integrator^31^ to maintain the secondary and tertiary structures of the protein (see SM Section I for further details on the sequence implementation). The key difference between the Z and M variants lies in the substitution of glutamic acid to lysine (E342K mutation) which reportedly delays the correct folding of the C-terminal disordered region (Fig. 1(b)). We first perform Direct Coexistence (DC) simulations^39,40^, in which a condensed and diluted protein phases cohabit within an elongated box. We employ DC simulations to compute the phase diagram in the temperature–density plane (see further details on DC calculations in Section SIII of the SM). Through this approach, we first determine the stability limits of both *α*_1_-antitrypsin variants. We ensure that our DC simulations do not present significant finite size effects by establishing a cross-section simulation box length which is ∼10 times larger than *α*_1_-antitrypsin radius of gyration, thus avoiding protein self-interactions through periodic boundary images^41–44^..

In Figure 1(c), the coexistence lines of both *α*_1_-antitrypsin sequences are shown, demonstrating the greater ability of Z-*α*_1_-antitrypsin to undergo LLPS and form protein condensates. We also include in Figure 1(c) rendered images of DC simulations for the M and Z variants at a temperature just above the critical one (T_*c*_) for the M variant, showing that the native form (Top panel) cannot form phase-separated condensates under these conditions while the Z variant (Bottom panel) undergoes LLPS. Importantly, higher critical solution temperatures have been proven to be associated with lower saturation concentrations in multiple *in silico* and *in vitro* studies of protein LLPS^45–50^, which strongly indicates that Z-*α*_1_-antitrypsin will form condensates (which could later potentially transition into pathological solid-like assemblies) at conditions at which the wild-type M variant would remain soluble. Moreover, the higher critical solution temperature for Z-*α*_1_-antitrypsin is consistent with the fact that intrinsically disordered regions enhance the ability to form condensates in many other proteins^51–54^ such as hnRNPA1, TDP-43 or FUS, where their low-complexity domains are crucial for enabling phase-separation^51,55^ . In the case of *α*_1_-antitrypsin, the mutated Z variant contains a disordered region (e.g., the C-terminus), that unless correctly folded into an intra-protein *β*-sheet structure^17^, it promotes condensate formation under less favorable conditions (i.e., at higher temperatures or at lower protein saturation concentrations).

Lastly, we generate two C-terminally truncated Z-*α*_1_-antitrypsin variants, Z-K367^*^ and Z-E387^*^, previously described as somatic escape variants by Brzozowska *et al*.^59^. In such study, the authors determined that C-terminal truncation in variants of Z-A1AT prevent multimerization, highlighting the role of the C-terminal region in forming higher molecular-weight assemblies^59^. Similarly, we measure their phase-separation behavior under the same conditions used for the M and Z variants. In contrast to the Z variant, which forms condensates across a wide temperature range (Figure 1(c)), neither Z-K367^*^ nor Z-E387^*^ show any detectable phase-separation behaviour at these conditions. This lack of phase-separation in the truncated variants underscores the major importance of the disordered C-terminal region—present in the Z variant—in promoting LLPS, and is consistent with prior reports targeting the C-terminal domain for being responsible of inter-protein association. This finding further highlights the capacity of our computational approach to reproduce experimentally observed trends.

### B. The disordered C-terminus of the Z variant enables greater intermolecular connectivity

We now investigate the molecular origin behind the greater propensity of Z-*α*_1_-antitrypsin to form condensates by computing the contact frequency map among all possible residue-residue interactions across the sequence. From our calculations shown in Fig. 1(c), we perform an analysis of the frequency of pairwise intermolecular contacts established by both *α*_1_-antitrypsin variants. To that goal, we examine the condensates formed at T/T_*c*_ ≃ 0.97, temperature at which the density of the condensates formed by both variants is similar (see Figure 1(c)). We consider an ‘effective’ intermolecular contact when two amino acids are closer than 1.2*σ*_*ij*_, being *σ*_*ij*_ = (*σ*_*i*_+*σ*_*j*_)*/*2, where *σ*_*i*_ and *σ*_*j*_ represent the molecular diameter of the two residues involved in such interaction as proposed in Refs.^24,45^.

In Figure 2 we show the intermolecular contact maps for both variants, where darker colors depict higher frequencies of pairwise residue-residue interactions. The average frequency of intermolecular contacts is expressed as a percentage, where 100% refers to a contact which persists across all the sampled configurations. Both variants show a similar pattern of intermolecular interactions until the 335th residue, where the C-terminus starts. Interestingly, domains from the 30th residue to the 45th, from the 102nd to the 110th, and from the 193rd to the 200th, which are enriched in charged and polar amino acids and remain intrinsically disordered, establish more frequent interactions than the rest of the sequence which is mostly structured. This is a common feature of multi-domain phase-separating proteins that combine both disordered and structured domains (through secondary and tertiary interactions) as previously reported for hnRNPA1^57,60^ or TDP-43^62–64^. Remarkably, the major difference between the contact frequency maps is observed in the C-terminus of both variants, differentiated from the rest of the sequence with dashed lines (Figure 2). As detailed in Section III A, this region harbours the E342K substitution that defines the Z variant, which disrupts the proper insertion of strand 5 of *β*-sheet A (s5A) and thereby hinders the efficient intramolecular folding of the C-terminal segment. Such destabilization increases the conformational flexibility and exposure of residues within the C-terminal region, allowing it to sample unfolded-like conformations. As a consequence, these residues can engage more frequently in transient intermolecular contacts with different domains from other *α*_1_-antitrypsin molecules. Our contact map analysis shows that these interactions occur predominantly between C-terminal regions, consistent with their increased structural lability in the Z variant. This behavior contrasts with the M variant, in which the C-terminus remains stably folded and largely unavailable for intermolecular interactions. Collectively, these results highlight the Z variant C-terminal region as a key driver of multivalent interactions, providing the molecular basis for its enhanced propensity to undergo multimerization.

**Figure 2:**
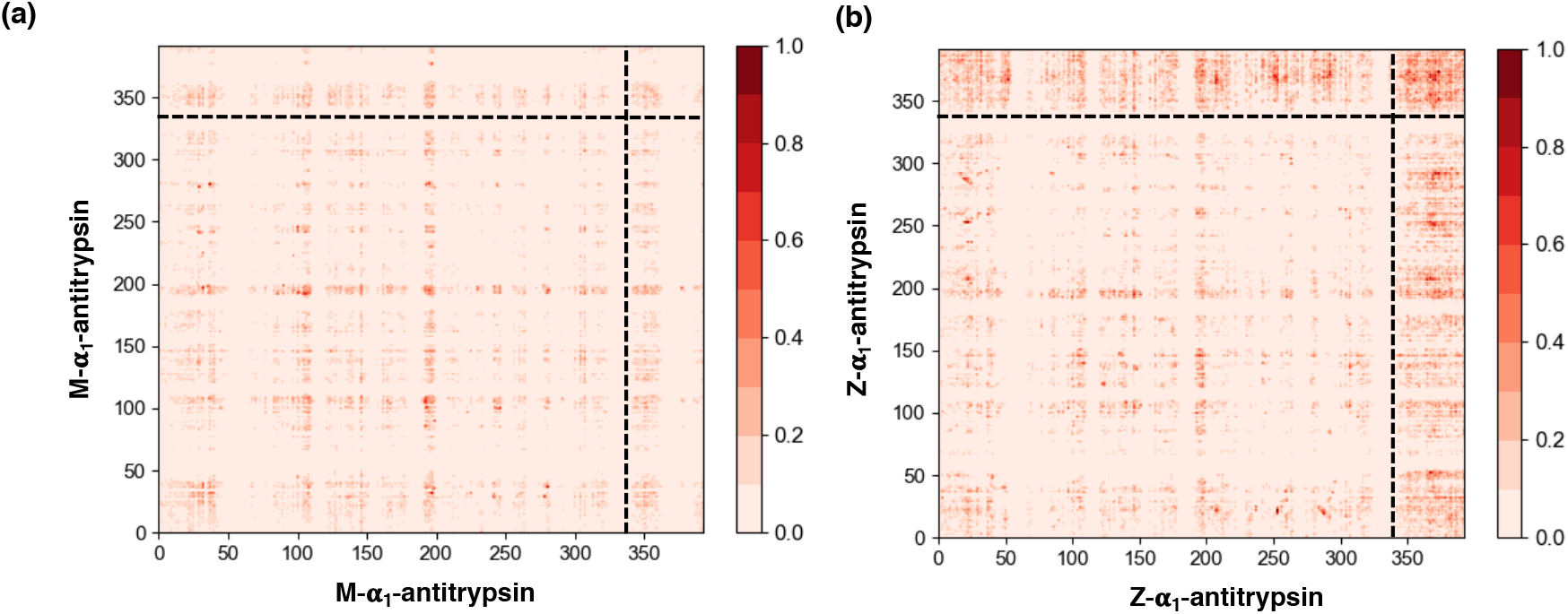
Average frequency of intermolecular contacts (expressed in %) per residue for (a) M-*α*_1_-antitrypsin and (b) Z-*α*_1_-antitrypsin. This analysis is performed over DC simulations (Figure 1(c)) at T/T_*c*_ ≃0.97. The black dashed line in both images indicates the region corresponding to the C-terminus, which remains disordered in the Z-*α*_1_-antitrypsin mutant, and folded in the M-*α*_1_-antitrypsin variant.

### C. Inter-protein *β*-sheet transitions promote condensate hardening

High-density local concentrations of disordered domains within protein condensates have previously been shown to induce the emergence of inter-protein *β*-sheets via structural transitions of low-complexity aromatic rich kinked segments^20,24,25,32,65^. While the folding of the M variant leads to the formation of intramolecular *β*-sheet structures, the disordered C-terminus of the Z intermediate variant has the potential to establish both intra- and inter-protein *β*-sheets within a protein condensed phase. Notably, a recent NMR study by Lowen et al.^66^ proposed that intermolecular *β*-sheet linkage between misfolded Z-*α*_1_-antitrypsin molecules can trigger an intramolecular conformational change in the acceptor molecule, causing a rearrangement that relocates the C-terminus to the opposite end of the molecule, exposing it for further intermolecular linkage with another misfolded serpin, thereby propagating the formation of cross-linked assemblies. This observation aligns with our proposed mechanism of extensive inter-molecular *β*-sheet formation, suggesting that such conformational transformations could underlie the transition from initially liquid-like condensed phases to solid-like assemblies.

Hence, after confirming the greater propensity of Z-*α*_1_-antitrypsin to undergo LLPS compared to its native form (Fig. 1(c)), we now perform non-equilibrium simulations (using our dynamic algorithm^23–25^; see SM Section II for further technical details on the algorithm), so that inter-protein *β*-sheets in the relevant sequence segments (dashed arrows in Figure 1(a)) can be formed either at intramolecular level (i.e., accomplishing the correct folding of the protein), or between different protein replicas establishing inter-protein *β*-sheets. Through the original dynamic algorithm^23,67^, we approximate *β*-sheet formation via disorder-to-order structural transitions by considering the atomistic implications (i.e., non-conservative strengthening of inter-protein binding, local protein rigidification, and changes in the intermolecular organization;^20,25^ ) of the gradual and irreversible accumulation of *β*-sheet structures in a time-dependent manner and as a function of the local protein density. Here, we adapt our dynamic algorithm^23,24^ to describe both intra- and inter-protein *β*-sheet transitions within condensates for evaluating the interplay between both folding phenomena. We perform these calculations at T/T_*c*_ =1.02, conditions at which M-*α*_1_-antitrypsin does not undergo LLPS, and hence, only condensate formation of Z-*α*_1_-antitrypsin occurs as experimentally found^17^.

In Figure 3(a) we plot the total number of *β*-sheets formed, distinguishing between those formed within the same protein (i.e., at intramolecular level; black solid line) and between different protein replicas (at inter-molecular level; red solid line). Strikingly, we observe a significantly larger amount of the latter, giving rise to local high-density protein clusters in which the different proteins are strongly engaged to each other through cross-*β*-sheet structures (as illustrated in Figure 3(b), including a zoom-in of a 3 protein cross-*β*-sheet cluster). In Figure 3(b)(right) we show a primitive path analysis (PPA)^24,25^ of the formed cross-*β*-sheet network to visualize the intermolecular connectivity across the condensate. For this calculation, we enforce that: (1) *β*-sheet domains formed across the C-terminus are fixed in space; (2) the intramolecular excluded volume is set to zero; and (3) the bond interaction has an equilibrium bond length of 0 nm. In this manner, the PPA algorithm minimizes the contour length of the protein strands that connect the different *β*-sheet domains in the C-terminal region, while preserving the topology of the underlying inter-protein network^68^, and allows for visualization of the connectivity generated by *β*-sheet clusters, which are mostly inter-protein arrays. Overall, our non-equilibrium simulations predict the ability of the Z-*α*_1_-antitrypsin variant to form phase-separated condensates, and subsequently transition into highly insoluble states through the stabilization of cross-*β*-sheet structures. Importantly, our simulations are performed under conditions at which the native form, M-*α*_1_-antitrypsin, does not form condensates, further validating our hypothesis since M-*α*_1_-antitrypsin has not been found in aberrant solid-like assemblies of AATD patients^14,69^.

**Figure 3:**
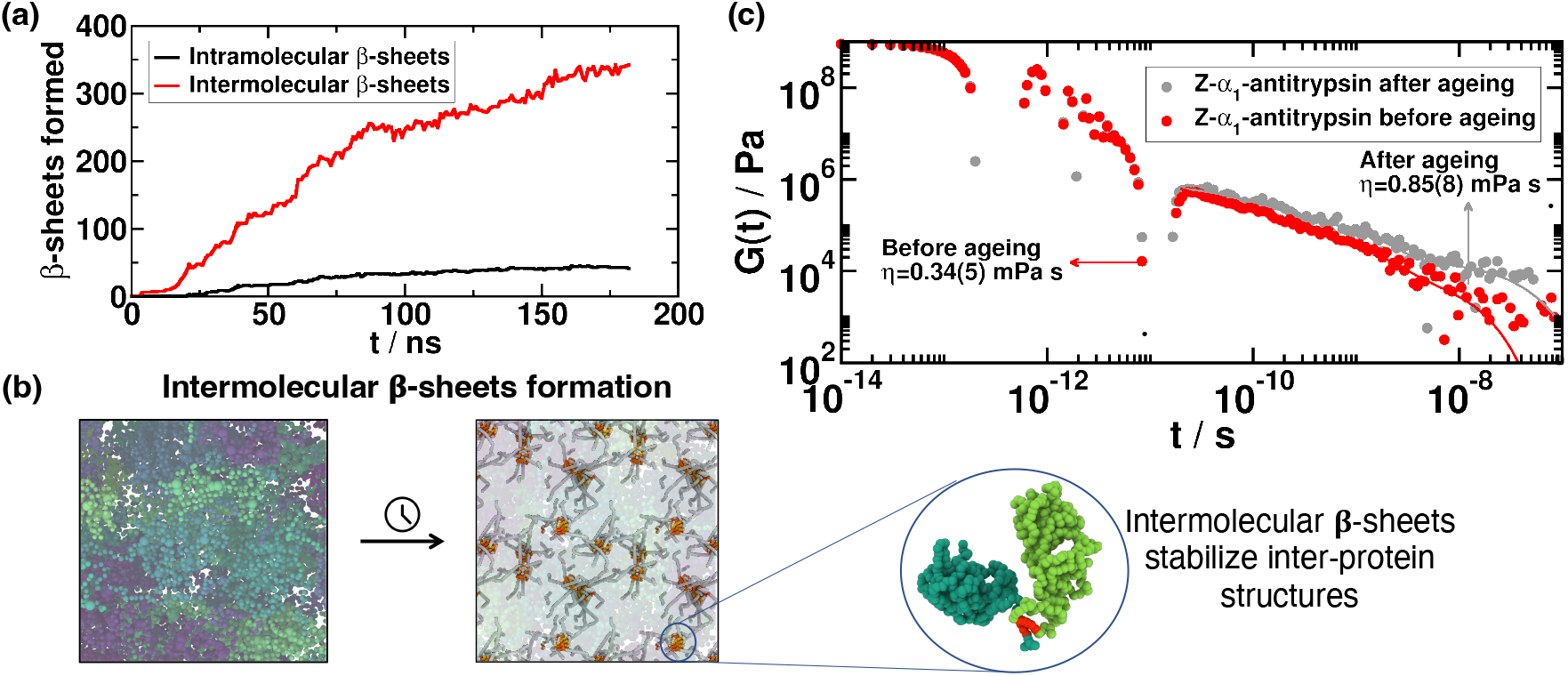
(a) Time-evolution of the different *β*-sheet structures formed (at inter- and intramolecular level) in Z-*α*_1_-antitrypsin condensates at T/T_*c*_ = 1.02 (referred T_*c*_ to that of the M-*α*_1_-antitrypsin variant) and the corresponding condensate equilibrium density at such temperature. (b) Primitive path analysis calculations showing the formation of inter-protein *β*-sheet structures among the C-terminus of the Z-*α*_1_-antitrypsin variant in phase-separated condensates. The zoomed-in image depicts a 2-protein cluster (dimer), stabilized through inter-protein *β*-sheet stacking. (c) Shear stress relaxation modulus (*G*(*t*)) of the Z-*α*_1_-antitrypsin bulk condensed phase at T/T_*c*_=1.02, prior to the formation of any inter-protein structure (red points), and after condensate maturation where inter-protein *β*-sheets are formed (grey points). Continuous lines depict Maxwell mode fits to *G*(*t*) data. From each set of points we obtain the viscosity for both systems (before and after condensate incubation time), by integrating in time the stress relaxation modulus. The error in the viscosity is estimated with the standard deviation when calculating this quantity using a different amount of Maxwell modes in the fit (see Section SIV).

Condensate maturation can lead to high viscous liquids or even solid-like states as the proteins within the condensates lose partial motility over time^51,70,71^. Such kinetically arrested dynamics is often associated with the emergence of condensate dysfunctional behavior, and has been widely linked to multiple neurological disorders^72,73^. Thus, we measure the viscosity of Z-*α*_1_-antitrypsin condensates (just after their formation, and after an incubation time of 200 ns) in order to assess the change in the droplet’s material properties over time due to the formation of inter-protein *β*-sheet structures. We note that the coarse-grained nature of our implicit-solvent model significantly underestimates the relaxation timescale of the proteins, and hence the viscosity of the condensates^45^; however, the observed trends and relative differences in the viscoelastic properties between initially formed and incubated condensates are expected to hold despite the artificially faster dynamics of our residue-resolution simulations^24^. We compute the shear stress relaxation modulus (*G*(*t*)) in Z-*α*_1_-antitrypsin bulk condensates prior (red curve) and post-maturation (gray curve in Fig. 3(c)) using separate NVT simulations, where we compute the auto-correlation function of any of the off-diagonal components of the pressure tensor^74,75^. Since the system is isotropic, an accurate expression of *G*(*t*) can be obtained by using the six independent components of the pressure tensor^76^ (see section SIV in SM). The viscosity of the system is then computed by integrating the stress relaxation modulus along time, using the Green-Kubo formula^77^

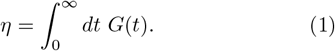

While at shorter timescales, *G*(*t*) easily converges and can be integrated numerically, for longer timescales (*t >* 10^−10^ s, Figure 3(c)) *G*(*t*) needs to be fitted to a series of Maxwell modes (*G*_*i*_ exp( −*t/τ*_*i*_)) equidistant in logarithmic time^74^, resulting in a function that is integrated analytically to obtain the viscosity (see Ref.^75^ or SM Section SIV for further details on these calculations).

In Figure 3(c), we show *G*(*t*) as a function of time along with the fits to the Maxwell modes at long timescales (solid lines). From these curves, we obtain the viscosity, being 0.336 mPa*·* s for Z-*α*_1_-antitrypsin before the formation of any inter-protein *β*-sheet structure, and 0.850 mPa *·*s for the incubated condensate. The shift in viscosity corresponds to the condensate maturation process of Z-*α*_1_-antitrypsin, which gradually transforms into a high-viscous phase which is locally arrested and stabilized by inter-protein *β*-sheet clusters (Fig. 3(b)). The emergence of Z-*α*_1_-antitrypsin foci has been consistently observed *in vivo* across multiple experimental studies^11,13,14,17^. In that respect, our simulations propose a mechanism that explains the experimentally observed formation of pathological Z-*α*_1_-antitrypsin assemblies through: (1) condensate formation via LLPS; and (2) progressive accumulation of inter-protein *β*-sheets that, when forming an interconnected percolated network (Figure 3(b)), increase the viscosity of the condensates over time (Figure 3(c)) and prevents their spontaneous dissolution.

## D. Heterogeneous nucleation of Z-*α*_1_-antitrypsin condensates

Nucleation of Z-*α*_1_-antitrypsin inclusion bodies naturally occurs within the confines of the endoplasmic reticulum (ER) membrane^17^. This observation suggests a possible mechanism by which the ER promotes heterogeneous condensate nucleation of Z-*α*_1_-antitrypsin over its surface. In this section, we investigate condensate formation near a flat structureless surface, and therefore elucidate the effect of an external barrier on the regulation of *α*_1_-antitrypsin LLPS.

As the specific intermolecular interactions responsible for protein self-assembly around a membrane are extremely challenging to describe with coarse-grained force fields^78,79^, we adopt two different modeling approaches for mimicking the ER surface. First, we introduce a non-specific interaction between a flat structureless wall and *α*_1_-antitrypsin using a soft attractive potential (see SM Section V for further details on the implementation). Essentially, the surface consists of a Lennard-Jones potential (with *ϵ*= 0.1 kcal mol^−1^ and *σ*= 15Å) inserted at the periphery of the simulation box (as illustrated in Figure 4(a)) which establishes a mild attractive region (of *<*0.5 k_*B*_T) that mimics non-specific chemical adsorption onto the ER membrane. Conversely, we model a repulsive surface by employing a purely repulsive interaction described by a harmonic potential *E* = *K*_*H*_ (*r* − *r*_*c*_)^2^ for *r < r*_*c*_. For the repulsive potential, the spring constant is set to *K*_*H*_ = 100 kcal mol^−1^ Å^−2^ and *r*_*c*_ = 4Å (see section SV of SM for further details). This results in a short-range moderately repulsive structureless surface located at the edge of the simulation box, in an analogous arrangement to the attractive barrier depicted in Figure 4(a). We note that these two surfaces represent oversimplified types of membrane-like interactions which cannot capture the biochemical complexity of the binding modes within an ER membrane. Nevertheless, they allow us to approximate generic physicochemical effects of confinement and surface proximity on protein condensation, which are the key aspects relevant to our study.

**Figure 4:**
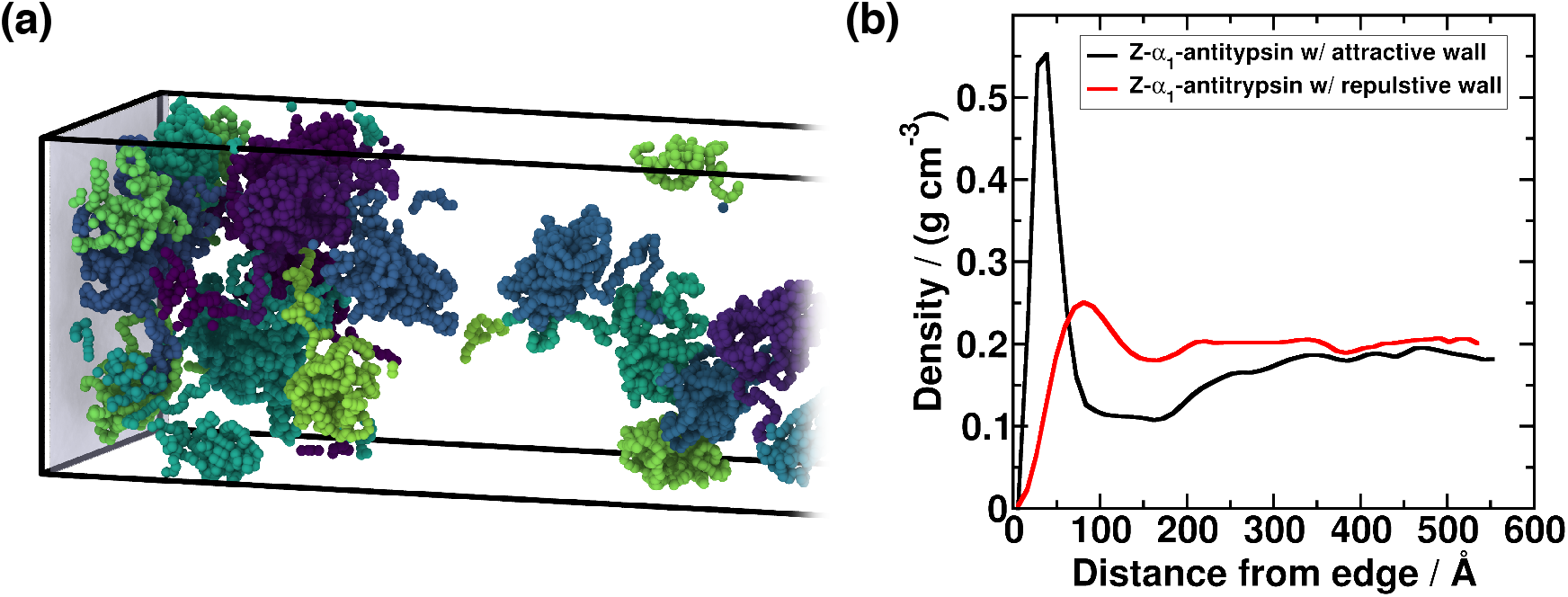
(a) Snapshot of a DC simulation of Z-*α*_1_-antitrypsin at *T/T*_*c*_ = 1.1 (where *T*_*c*_ corresponds to the critical temperature for LLPS of the Z variant) in presence of an external surface indicated by the shaded plane at the edge of the simulation box. In the snapshot, each protein replica is represented in a different colour. (b) Averaged density profiles along the long axis of the DC simulation box, evaluated from the external surface. The nature of the inserted surface is indicated in the figure legend.

We perform DC simulations of the Z-*α*_1_-antitrypsin variant well above its critical solution temperature (i.e., at *T/T*_*c*_ = 1.1 where *T*_*c*_ corresponds to the critical temperature for LLPS of the Z variant) in the presence of the moderately attractive and repulsive surfaces. At temperatures *T* ≥ *T*_*c*_, phase separation is unattainable, and therefore the emergence of condensates would only be a direct consequence of the presence of an external surface. We note that, due to the calculation setup, the simulation is not periodic along the elongated axis, as the surface remains uncrossable by the molecules. In Figure 4(b), we present density profiles along the long axis of the simulation box, starting from the edge where the attractive (or repulsive) surface is located. For the attractive surface (black curve), we observe a local density maximum induced by the external layer. Despite using a weak attractive interaction (i.e., lower than 0.5 k_*B*_T), this density maximum indicates the presence of protein clusters with densities that correspond to those of condensates well below the critical solution temperature (please note that the critical density is ∼0.2 g/cm^3^; Fig. 1(c)). These clusters exhibit significant local high-density fluctuations, which over time can induce the formation of inter-protein *β*-sheets, further increasing their stability and viscosity, and originating the characteristic features of insoluble condensates as described in Section III C and Fig. 3.

Additionally, for the simulation with a repulsive surface (depicted in Fig. 4(b) by the red curve), we observe a density maximum near the external layer, although less pronounced than for the moderately attractive wall (black curve). Although this maximum does not reach average densities exceeding 0.25 g cm^−3^, our results suggest a significant propensity of Z-*α*_1_-antitrypsin to form clusters on an external surface, despite lacking an explicit physicochemical affinity for the wall (i.e., attractive cross-interactions). This behavior can be attributed to the lower interfacial free energy that proteins may exhibit with the wall, compared to the surface tension between the condensate and the protein diluted phase. Under such circumstances, condensates would form on the surface through a classical heterogeneous nucleation mechanism^80,81^. In Ref.^82^, a similar mechanism was reported for hard-sphere colloids in the presence of purely repulsive, flat structureless walls, which favor the nucleation of crystalline clusters from the supersaturated fluid, leading to much higher nucleation rates compared to those observed in the absence of surfaces (i.e., homogeneous crystallization^83^). This phenomenon is driven by the lower wall-nucleus interfacial free energy promoting cluster wetting on the surface^84^, and appears to be analogous to the behavior observed in our simulations for Z-*α*_1_-antitrypsin. Remarkably, even in the case of the purely repulsive wall, the reported density maximum is of the order of condensate densities below the critical solution temperature (see phase diagram in Fig. 1(c)). Hence, protein surface coating may induce local high-density clusters that in turn promote inter-protein *β*-sheet transitions of Z-*α*_1_-antitrypsin at conditions where LLPS is not expected to spontaneously occur, and subsequently trigger the formation of highly viscous condensates (Fig. 3). Overall, the results of Figure 4 reflect how the presence of either attractive or repulsive walls, enhances Z-*α*_1_-antitrypsin propensity to form condensates, matching experimental observations from Ref.^17^ where aberrant solid-like assemblies appear attached to ER membranes.

## IV. DISCUSSION

The results presented in this work shed light on the intricate phase behavior of the misfolded Z-*α*_1_-antitrypsin variant and its implications in the development of AATD^11,13^ . Through computational molecular modeling, we have elucidated the impact of the structural differences between the native M-*α*_1_-antitrypsin sequence and the misfolded Z variant, particularly focusing on the phase behavior of the protein depending on the folding state of its C-terminus. The misfolded intermediate of Z-*α*_1_-antitrypsin exhibits an intrinsically disordered C-terminus, contrasting with the structured intramolecular *β*-sheet domain found in the native M variant (Fig. 1(a) and 1(b)). We examine their phase diagram by means of Direct Coexistence simulations finding a greater propensity of the Z variant to undergo LLPS compared to the M native sequence (Fig. 1(c)). These results are consistent with previous experimental findings^14,17^, and further validate our coarse-grained model in describing their coexistence lines^26^. The increased propensity for phase separation is unambiguously attributed to the presence of an intrinsically disordered region in the Z variant which is capable of establishing multiple intermolecular contacts with neighboring C-terminal domains of adjacent protein replicas (Figure 2(b)). Moreover, the disordered domain also facilitates intermolecular interactions with most regions of the sequence compared to the M variant in which such domain is practically inert. The enhanced ability of the Z variant to form condensates favors the progressive formation of inter-protein *β*-sheet clusters instead of individual intramolecular folding. Consequently, the emergence of interconnected strong-binding networks via inter-protein *β*-sheet accumulation significantly stabilizes the formation of condensates against their spontaneous dissolution.

We measure the viscosity of Z-*α*_1_-antitrypsin condensates once they are formed (i.e., in the absence of inter-protein *β*-sheet clusters) and after incubation. The progressive emergence of cross-*β*-sheet clusters leads to an increase of condensate viscosity due to the inter-protein assembly of local ordered structures that mutually bind via strongly *π*-*π* interactions and hydrogen bonding between backbone interactions^34^. These highly stable cross-*β*-sheet structures^25,32^ lead to the progressive hardening of initially liquid-like condensates. The increased viscosity observed in aged Z-*α*_1_-antitrypsin condensates provides a mechanistic explanation for the aberrant formation of misfolded Z-*α*_1_-antitrypsin inclusion bodies found in vivo^11,13,14,17^.

Furthermore, experimental evidence^17^ suggests that the formation of aberrant Z variant clusters in the ER membrane may be promoted under confinement conditions, or nucleated over its surface. We carry out simulations of Z-*α*_1_-antitrypsin in presence of both repulsive and moderately attractive flat structureless walls to elucidate their impact on protein phase-separation. Strikingly, we observe that both types of external surfaces (independently of their chemical adsorption affinity) induce local high-density fluctuations above the characteristic coexistence condensate densities to undergo LLPS (Fig. 4(b)). This suggests a heterogeneous nucleation mechanism by which Z-*α*_1_-antitrypsin (or M-*α*_1_-antitrypsin under higher concentrations) aided by an external surface could potentially achieve the critical saturation concentration to form small condensate nuclei. The initial nuclei can be further stabilized and grow through the formation of cross-*β*-sheet clusters which gradually increase the inter-protein interaction strength, and thus, the condensate viscosity and stability (Fig. 3(c)).

Altogether, our study advances our understanding of the intricate interplay between structure and phase behaviour in M- and Z-*α*_1_-antitrypsin, as well as on their role in driving AATD pathogenesis. The proposed mechanism to explain its aberrant condensation is based on the formation of phase-separated droplets stabilized by C-terminus intermolecular interactions which over time develop inter-protein *β*-sheet transitions. This mechanism provides a strong hypothesis for understanding the onset of AATD. Furthermore, our findings at the molecular level can contribute to the development of therapeutic strategies^85^ targeting the protein interactions involved in the accumulation of *α*_1_-antitrypsin inclusion bodies at the ER. Potential routes could focus on the intramolecular folding of the C-terminus, or the selective engagement of external molecules with this domain to inhibit inter-protein *β*-sheet accumulation.

## ACKNOWLEDGMENTS

S.-B. acknowledges funding from the Derek Brewer scholarship of Emmanuel College and EPSRC Doctoral Training Programme studentship, number EP/T517847/1, Ramon y Cajal fellowship (awarded to J.R.E.), as well as the UKRI EPSRC under the UK Government’s guarantee scheme (EP/Z002028/1), following successful evaluation by the ERC (Consolidator Grant awarded to R.C.G.) under the European Union’s Horizon Europe research and innovation programme. R.C.-G. acknowledges funding from the European Research Council (ERC) under the European Union Horizon 2020 research and innovation programme (grant agreement 803326). A. T. is funded by European Research Council (ERC) under the European Union Horizon 2020 research and innovation programme (grant agreement 803326) and Ramon y Cajal fellowship (RYC2021-030937-I). J. R. E. also acknowledges funding from the Ramon y Cajal fellowship (RYC2021-030937-I) and the Spanish National Agency for Research (PID2022-136919NA-C33). This work has been performed using resources provided by the Cambridge Tier-2 system operated by the University of Cambridge Research Computing Service (http://www.hpc.cam.ac.uk) funded by EPSRC Tier-2 capital grant EP/P020259/1. The authors also thankfully acknowledge RES computational resources provided by MareNostrum 5 through the activities FI-2024-3-0001 and FI-2025-1-0008. The funders had no role in study design, data collection and analysis, decision to publish, or preparation of the manuscript.

## DATA AVAILABILITY

The data that supports the findings of this study are available within the article and its Supplementary Material. Moreover, example files for (i) Direct Coexistence simulations of the M variant (ii) Direct Coexistence simulations of the Z variant (iii) Non-equibrium simulations of the Z variant (iv) MD simulations of the Z variant in presence of an attractive wall (iv) MD simulations in presence of a repulsive wall; can be found in the Zenodo repository https://doi.org/10.5281/zenodo.17193274

## SUPPLEMENTARY MATERIAL LIST OF LEGENDS

No figures, tables, or movies are included. The Supporting Material consists of a detailed description of the CALVADOS2 model, the ageing algorithm, methodology regarding the calculation of phase diagrams and viscoelastic properties, and the surface-driven condensation simulation details.

