## Supplementary Material for "Molecular insights on the mechanism of *α*_1_-antitrypsin condensate formation and maturation"

### Supplementary Material: Molecular insights on the mechanism of $\alpha_1$-antitrypsin condensate formation and maturation


Ignacio Sanchez-Burgos
*Maxwell Centre, Cavendish Laboratory, Department of Physics, University of Cambridge, J J Thomson Avenue, Cambridge CB3 0HE, United Kingdom.*

Andres R. Tejedor
*Yusuf Hamied Department of Chemistry, University of Cambridge, Lensfield Road, Cambridge CB2 1EW, United Kingdom. and Department of Physical-Chemistry, Universidad Complutense de Madrid, Av. Complutense s/n, 28040, Madrid, Spain.*

Rosana Collepardo-Guevara
*Yusuf Hamied Department of Chemistry, University of Cambridge, Lensfield Road, Cambridge CB2 1EW, United Kingdom.*

Jorge Bernardino de la Serna
*Faculty of Medicine, Imperial College London, Ayrton Rd, South Kensington, London SW7 5NH, United Kingdom.*

Jorge R. Espinosa**
*Maxwell Centre, Cavendish Laboratory, Department of Physics, University of Cambridge, J J Thomson Avenue, Cambridge CB3 0HE, United Kingdom. and Department of Physical-Chemistry, Universidad Complutense de Madrid, Av. Complutense s/n, 28040, Madrid, Spain.*


(Dated: March 11, 2025)


#### SUPPLEMENTARY SECTION I. CALVADOS2 MODEL

In order to simulate $\alpha_1$-antitrypsin, we use the CALVADOS2 chemically-accurate coarse-grained (CG) protein model proposed by Tesei *et al.* [1] implemented in the Molecular Dynamics LAMMPS package [2]. The coarse-grained model resolution is of one bead per amino acid. In the model, the intrinsically disordered regions (IDRs) of the proteins are considered as fully flexible polymers, while the structured globular domains are treated as rigid bodies (where their conformations are taken from the Protein Data Bank (PDB) crystalline structure by using the rigid body integrator of LAMMPS [2]. Moreover, the interactions of the structured globular domains are scaled down by a 30% to account for the 'buried' amino acids as proposed by Krainer *et al.* [3].

The potential energy of the coarse-grained CALVADOS2 force field is given by:

$$E = E_{\text{Bonds}} + E_{\text{Electrostatic}} + E_{\text{Hydrophobic}}, \tag{S1}$$

where $E_{\text{Hydrophobic}}$ and $E_{\text{Electrostatic}}$ interactions are only applied between non-bonded beads and $E_{\text{Bonds}}$ between subsequent beads directly bonded to each other. Bonded interactions between subsequent amino acid protein beads or consecutive RNA nucleotides are described by an harmonic potential:

$$E_{\text{Bonds}} = \sum_{\text{Protein bonds}} k(r - r_0)^2, \tag{S2}$$

where the equilibrium bond length is $r_0 = 3.81\text{Å}$ between bonded amino acid beads. The spring constant is $k = 10\,\text{kJ}/(\text{mol}\text{Å}^2)$. The electrostatic interactions, $E_{\text{Electrostatic}}$, between charged amino acids are described by a Coulomb/Debye-Hückel potential of the form:


*

$$E_{\text{Electrostatic}} = \sum_i \sum_{j<i} \frac{1}{4\pi D} \frac{q_i q_j}{r} e^{-r/\kappa}, \tag{S3}$$

where $q_i$ and $q_j$ represent the charges of the beads $i$ and $j$. Within this model the Debye length ($\kappa$) is not a fixed value, but rather depends on the salt concentration, allowing for a better description of LLPS modulation with varying salt concentration. The Debye length is defined as:

$$\kappa = \sqrt{\frac{1}{8\pi B c_s}} \tag{S4}$$

where $c_s$ is the ionic strength and $B$ is the Bjerrum length, defined as:

$$B = \frac{1.671x10^{-5}}{DT}. \tag{S5}$$

In this model, the relative dielectric constant of the solvent ($D$) follows the empirical equation [4]:

$$D = \frac{5321}{T} + 233.76 - 0.9297T + 1.417x10^{-3}T^2 - 8.292x10^{-7}T^3 \tag{S6}$$

The hydrophobic interactions between different amino acid types are described by:

$$E_{\text{Hydrophobic}} = \sum_i \sum_{j<i} \begin{cases} 4\epsilon_{ij}\left[\left(\frac{\sigma_{ij}}{r}\right)^{12} - \left(\frac{\sigma_{ij}}{r}\right)^{6}\right] + (1-\lambda_{ij})\epsilon_{ij}, & r < 2^{1/6}\sigma_{ij} \\ \lambda_{ij} 4\epsilon_{ij}\left[\left(\frac{\sigma_{ij}}{r}\right)^{12} - \left(\frac{\sigma_{ij}}{r}\right)^{6}\right], & \text{otherwise,} \end{cases} \tag{S7}$$

where $\lambda_i$ and $\lambda_j$ are parameters that account for the hydrophobicity of the $i$th and $j$th interacting particles respectively, being $\lambda_{ij} = (\lambda_i + \lambda_j)/2$. The excluded volume of the different residues/nucleotides is given by $\sigma_i$ and $\sigma_j$, where $\sigma_{ij} = (\sigma_i + \sigma_j)/2$, and $r$ is the distance between the $ij$ particles. $\epsilon_{ij}$ is set to 0.2 kcal/mol. When at least one of the $ij$ amino acids is part of a structured globular domain, $\lambda_{ij}$ is scaled by a factor of 0.7 to account for the 'buried' amino acids in globular domains [3]. The specific values for each amino acid and nucleotide $\sigma$, $q$, and $\lambda$ parameters can be found in reference [1].

#### SUPPLEMENTARY SECTION II. AGEING ALGORITHM

To dynamically mimic the structural diversity of proteins during condensate maturation, we implement a time-dependent and local-dependent algorithm that modulates the strength of disordered *vs.* structured interactions. Low-complexity aromatic-rich kinked segments (LARKS) can transition from their fully disordered state to form inter-peptide structured $\beta$-sheets depending on the local environment. Every 100 simulation timesteps, this dynamical algorithm evaluates whether the conditions around each fully disordered LARKS across the protein sequences are favorable for undergoing an 'effective' disorder-to-order cross-$\beta$-sheet transition, and thus, modifies the interaction parameter $\lambda$ (Eq. S7) to those corresponding to inter-peptide structured $\beta$-sheet motifs. We make use of a distance criterion, so that if 4 peptides are within a certain cut-off distance, the average effective interaction in the aforementioned becomes four times stronger, in line with our previous work im which this interaction was determined through atomistic Potential of Mean Force calculations for other proteins [5–8]. Additionally, to mimic the higher rigidity of inter-peptide $\beta$-sheet motifs upon a structural transition, we introduce an angular term to the total energy (equation S1) that follows the next equation:

$$E_{\text{Angles}} = k_{ang}(\theta - \theta_0)^2, \tag{S8}$$

where we set $\theta_0 = 180°$ and $k_{ang} = 5\ kcal\ mol^{-1}\ rad^{-2}$. To carry out these non-equilibrium simulations, we used the USER-REACTION [9] package of LAMMPS which allows to change the topology of the molecules on a time-dependent and local-dependent manner across the simulations.

The criterion that at least four peptides should be in close contact to trigger a disorder-to-order transition has been chosen based on the following arguments. Four interacting peptides is the minimal system where all the different types of stacking and hydrogen bonding interactions that stabilize the $\beta$-sheet fibrillar ladder are fulfilled; i.e., two interacting steps of the ladder each made of a pair of $\beta$-sheet peptides [5]. Thus, considering fewer interacting peptides (e.g. only three or two) would severely underestimate the strength of interactions among ordered LARKS, and subsequently, erroneously preclude the formation of kinetically arrested states at the coarse-grained level. On the other hand, if we made the criterion even more stringent (i.e., by requiring clustering of five or more peptides), the strength of interactions among the system would remain consistent with kinetic arrest at the coarse-grained level. However, such criterion would now render the coarse-grained simulations prohibitively expensive.

#### SUPPLEMENTARY SECTION III. OBTAINING PHASE DIAGRAMS VIA DIRECT COEXISTENCE SIMULATIONS

To calculate the coexisting densities of the phase diagrams, we employ the Direct Coexistence method [10–12]. Within this scheme, the two coexisting phases are simulated by preparing periodically extended slabs of the two phases, the condensed and the diluted phase, in the same simulation box. Once our DC simulations have reached equilibrium, we compute the density profile along the long axis of the box, and thus, we extract the density of the two coexisting phases. To estimate the critical point of the phase diagrams, we use the universal scaling law of coexistence densities near a critical point [13], and the law of rectilinear diameters [14]:

$$\big(\rho_l(T) - \rho_v(T)\big)^{3.06} = d\left(1 - \frac{T}{T_c}\right) \tag{S9}$$

and

$$(\rho_l(T) + \rho_v(T))/2 = \rho_c + s_2(T_c - T) \tag{S10}$$

where $\rho_l$ and $\rho_v$ refer to the coexisting densities of the condensed and diluted phases respectively, $\rho_c$ is the critical density, $T_c$ is the critical temperature, and $d$ and $s_2$ are fitting parameters.

#### SUPPLEMENTARY SECTION IV. VISCOSITY CALCULATIONS

From independent bulk simulations, we can compute the viscosity in the condensate. The shear viscosity can be straightforwardly calculated by integrating the relaxation modulus in time (see Chapter 7 of the book [15]):

$$\eta = \int_0^\infty dt\, G(t) \tag{S11}$$

In an isotropic system, we can compute the shear relaxation modulus $G(t)$ more accurately by using all the components of the pressure tensor ($\sigma_{\alpha\beta}$) as shown in Ref. [16]:

$$\begin{aligned} G(t) = {} & \frac{V}{5k_BT}\left[\langle\sigma_{xy}(0)\sigma_{xy}(t)\rangle + \langle\sigma_{xz}(0)\sigma_{xz}(t)\rangle + \langle\sigma_{yz}(0)\sigma_{yz}(t)\rangle\right] \\ & + \frac{V}{30k_BT}\left[\langle N_{xy}(0)N_{xy}(t)\rangle + \langle N_{xz}(0)N_{xz}(t)\rangle + \langle N_{yz}(0)N_{yz}(t)\rangle\right], \end{aligned} \tag{S12}$$

where $N_{\alpha\beta} = \sigma_{\alpha\alpha} - \sigma_{\beta\beta}$ is the first normal stress difference. This correlation can be easily computed by using the compute ave/correlate/long in the USER-MISC package of LAMMPS [2]. In all cases, the relaxation modulus presents an initial regime that mainly accounts for the intramolecular interactions, followed by a terminal region which corresponds to much slower relaxation modes, as those coming from intermolecular interactions and the relaxation of the large scale protein and RNA conformations. Due to the very wide range of timescales involved in the calculation and the noisy nature of the relaxation modulus in the terminal region obtained in the simulations, we follow a particular strategy to calculate our estimate of viscosity. At short times, $G(t)$ is smooth and the integral can be computed using numerical integration (trapezoidal rule). However, at longer times $G(t)$ presents more noise, and hence, we calculate the integral in that regime by first fitting $G(t)$ to a series of Maxwell modes ($G_i \exp(-t/\tau)$) equidistant in logarithmic time [17] and then by calculating the integral analytically. Our fit to the Maxwell modes is carried out with the help of the open-source RepTate software (version 1.1.1 20200602) [18]. Finally, viscosity is obtained by adding the two terms:

$$\eta = \eta(t_0) + \int_{t_0}^{\infty} dt\, G_M(t), \tag{S13}$$

where $\eta(t_0)$ corresponds to the computed term for short time-scales, $G_M(t)$ is the part evaluated via the Maxwell modes fit at long time-scales, and $t_0$ is the time that separates both.

#### SUPPLEMENTARY SECTION V. SURFACE-DRIVEN CONDENSATION SIMULATIONS

As mentioned in the main text, we use two modelling approaches to mimic the presence of a non-specific surface in our simulations. This is done by inserting an additional term to the potential energy. In this way, the system does not present periodic boundary conditions in the direction perpendicular to the inserted surface. More specifically, we insert an dispersive and a purely repulsive potentials (in separate simulations). For the dispersive surface, we make use of the *fix wall/lj126* command in LAMMPS [2], which adds a surface of Lennard-Jones interaction:

$$E_{wall\ LJ} = 4\epsilon \left[ \left(\frac{\sigma}{r}\right)^{12} - \left(\frac{\sigma}{r}\right)^{6} \right] \quad if\ r < r_c, \tag{S14}$$

where $r$ is the distance separating the surface and the amino acids, $r_c$ is the cut-off distance for the interaction, $\epsilon = 0.1$ kcal/mol, $\sigma = 15$ Å and $r_c = 37.5$ Å. For the case of the repulsive surface, we use the *fix wall/harmonic command*, which introduces a repulsive-only harmonic spring potential:

$$E_{wall\ repulsive} = K_H (r - r_c)^2 \quad if\ r < r_c, \tag{S15}$$

where we set $K_H = 100$ kcal mol$^{-1}$ Å$^{-2}$ and $r_c = 4$ Å. Note that the surface matches the boundary of the simulation box, and is applied in both (positive and negative) directions.

#### SUPPLEMENTARY SECTION VI. REFERENCES
